# Evolution of reduced dormancy during range expansions

**DOI:** 10.1101/2021.10.11.463894

**Authors:** Justin M. J. Travis, Poppy Mynard, Greta Bocedi

## Abstract

There is increasing evidence that life-history traits can evolve rapidly during range expansion and that this evolution can impact the ecological dynamics of population spread. While dispersal evolution during range expansion has received substantial attention, dormancy (dispersal in time) has not. Here, we use an individual-based model to investigate the evolution of seed dormancy during range expansion. When a population is at spatial equilibrium our model produces results that are consistent with previous theoretical studies: seed dormancy evolves due to kin competition and the degree of dormancy increases as temporal environmental variation increases. During range expansions we consistently observe evolution towards reduced rates of dormancy at the front. Behind the front there is selection for higher rates of dormancy. Notably, the decreased dormancy towards the expanding margin reduces the regional resilience of recently expanded populations to a series of harsh years. We discuss how dormancy evolution during range expansion, and its consequences for spatial population dynamics, may impact other evolutionary responses to environmental change. We end with suggestions for future theoretical and empirical work.

## Introduction

Over the last two decades there has been substantial attention devoted to the role that evolutionary processes can play during range expansions [1], both in invasive species (see reviews by [2–4] and in species expanding their ranges into newly suitable regions under climate change [5–9]. Hybridisation (e.g., [10,11], local adaptation (e.g. [12–15]) and life history evolution (e.g., selfing rates [16]; resistance to herbivores [17]; dispersal behaviour [18,19]) have all been implicated as determinants of either the probability that an introduction leads to an invasion, or the spatial dynamics of the invasion. Similarly, for species shifting their ranges due to climate change, local adaptation and the evolution of a range of life-history traits including dispersal and mating systems (e.g., [6–8]) have been highlighted as having potentially substantial impacts on the range dynamics.

In both the invasion literature and the range shifting literature, there has been a major focus on the role that dispersal evolution can play in driving the dynamics of population spread. In an early model Travis & Dytham [20] demonstrated that range expansion may be accelerated as greater rates of dispersal will generally evolve. However, their work showed that when even a weak Allee effect was incorporated, the evolution of increased dispersal was substantially reduced. Further theory, considering the evolution of density-dependent dispersal, has demonstrated that during range expansions selection favours strategies that yield moderate rates of emigration even from patches where density (and intraspecific competition) is very low [21]. Other work has highlighted the distinctiveness of the evolutionary process during range expansion, drawing attention to what has been dubbed spatial sorting [22,23]. In the context of dispersal evolution, high dispersal phenotypes are sorted at the expanding front and are thus likely to reproduce with one another, potentially resulting in individuals of even greater dispersal propensity and ability. In models of species shifting their ranges due to climate change, dispersal evolution can result in elastic margins [5,24]. Elevated dispersal at the expansion front supports sink populations in highly marginal habitat for a transient period after climate change ends and before selection operates to reduce dispersal back to the level expected at a stationary range margin (at which point there is less dispersal to prop up sink populations). Alongside the development of theory, there is increasing empirical evidence for rapid dispersal evolution during range expansions [25–28]. This comes from a range of taxa, for invasions and climate-induced range shifts, and in both natural settings and in experiments. The now classic example for an invasive species is the cane toad invading Australia [19,29,30]. Selection has resulted in substantial changes to multiple dispersal traits that together have resulted in the species now range expanding at least 5 times as rapidly now as in the earlier phase of expansion.

While the eco-evolutionary dynamics of dispersal have been well-studied in the context of range expansion, there has been little consideration of the role for what has often been termed ‘dispersal in time’ – i.e., dormancy [31,32]. This is surprising given both how widely dormancy is exhibited within animals, plants and microbes and the substantial role that it can play in population and community dynamics. We lack theoretical predictions into how dormancy should evolve during range expansions and on how this is likely to impact the ecological dynamics of population spread. Similarly, we lack empirical studies documenting potential changes in dormancy during either invasions or climate-driven range expansions.

While we lack studies investigating dormancy evolution during range expansions, there is a substantial literature focused on how dormancy evolves in stationary ranges. Theoretical studies have demonstrated that seed dormancy can evolve even in temporally stable environments [33–36], by reducing the number of sibling seeds germinating simultaneously. Heterogeneity in siblings’ dormancy rates reduces kin competition and increases a plant’s inclusive fitness [36,37]. However, it is in temporally variable environments where the strongest selection for seed dormancy is likely to occur [38–40]. Under these conditions dormancy can function as a bet hedging strategy (e.g. [41,42,42]): seed dormancy spreads the risk of germination over time, and is increasingly advantageous the greater the frequency of bad years [43].

In addition to work focused on understanding the drivers of dormancy evolution, there is strong evidence of a range of important consequences that dormancy can have on ecological dynamics. This includes the role that dormancy can play in increasing species diversity by enabling coexistence of competitors through the storage effect (e.g. [44,45]), and more generally evidence for the role of dormancy, together with dispersal, in structuring metacommunities [46]. Dormancy can also have substantial impacts on evolutionary processes [47]. The major role dormancy plays in ecological and evolutionary dynamics is not limited to eukaryotes and there has been substantial interest in understanding the causes and consequences of microbial dormancy over the last decade [48,49]. Furthermore, there is interest in considering the eco-evolutionary dynamics of dormancy in the context of cancers [50]. Given the major roles that dormancy can play in driving ecological and evolutionary outcomes across a broad range of systems, it is important that we gain understanding of how range expansions (of eukaryotes, prokaryotes, or even cancer cells) are likely to impact dormancy dynamics.

Here we build an individual-based model to investigate the eco-evolutionary dynamics of seed dormancy during range expansions. We run sets of simulations designed to address three key issues. First, we assume a fixed rate of dormancy and ask ***how dormancy influences the rate of a range expansion*** under both temporally stable and temporally variable environments. Second, we determine ***how dormancy evolves during range expansion***, and again consider how this differs under stable and variable environments. Third, we address the question of ***how dormancy evolution impacts the ecological dynamics of range expanding species***, focusing both on the rate of expansion and on the spatial dynamics in recently colonised regions.

## The Model

We develop an individual-based, spatially explicit simulation model to investigate the evolution of seed dormancy during range expansion. Simulations take place in an arena of cells (dimensions x = 400, y = 50), with each cell having a carrying capacity of *K* adult plants. We model an annual plant reproducing asexually. The model runs in discrete time and the ordering of events is as follows: seed production, seed dispersal, germination, seedlings density regulation. This formulation has much in common with similar models used to tackle a wide range of questions within evolutionary ecology (e.g., [51–54]).

Adult plants each produce *s* seeds. *s* can either be a constant for the duration of a model run (i.e., temporally constant environment), or can be affected by temporal environmental stochasticity and hence vary from year to year. In the latter case, *s* at a given year *t* is given by:

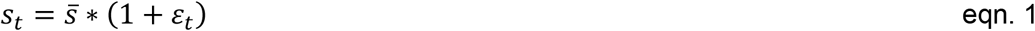

where 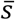 is the mean fecundity 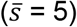 and *ε* is an environmental noise value generated from a first-order autoregressive process [55]:

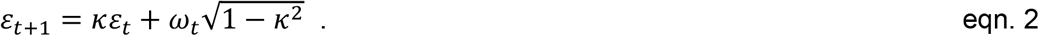

Here, *κ* is the autocorrelation coefficient and *ω* is a random normal variable drawn from N(0,*σ*), where *σ* changes the amplitude of the fluctuations. We assume temporally uncorrelated noise (i.e., white noise; *κ* = 0) and apply it uniformly across the landscape. Thus, seed production, *s*, is the same in every patch on the landscape in any given year. We further assume 0 ≤ *s* ≤ 10.

Each seed has a probability *d* = 0.1 of dispersing. Dispersing seeds move at random to one of their natal cell’s nearest eight neighbours. We wrap the landscape across the y-axis, while individual dispersing beyond the edges along the x-axis are lost. After dispersal, all seeds enter a seed bank. Every year, each seed germinates with probability 1 - *γ*, where *γ* is the probability of dormancy and can be either constant or determined by the seed’s genotype in case of dormancy evolution. Seeds that do not germinate remain in the seed bank with probability 1 - *m*, where *m* is the annual rate of mortality associated with remaining dormant. Following germination, seedling density is regulated (seedling competition), whereby the seedling survival probability is given by min(*K* / *N*_*seedling*_, 1), where *N*_*seedling*_ is the total number of seedlings present in the cell. All surviving seedlings are then developing to adults and reproduce the following year.

In the case of evolution of dormancy, every individual carries a quantitative character (single haploid gene with continuous alleles), *γ*, that codes for dormancy propensity. We assume asexual reproduction, and seeds inherit their genotype from their parent. There may be mutation events associated with seed production. Mutations to *γ* occur with probability *β* = 0.01/allele/year. When a mutation occurs, the individual genotype is altered by adding a value sampled from a uniform real distribution U(−0.1, 0.1). We constrain the genotype such that 0.0 ≤ *γ* ≤ 1.0.

Initially, we run a set of simulations to establish how a fixed rate of dormancy impacts range expansion dynamics. We do this both for a temporally stable environment and for a temporally variable environment (white noise). In both cases, we run the simulation twenty times for rates of seed dormancy between 0.0 and 0.95 in increments of 0.05. In each simulation, we introduce *K* = 25 individuals into a single patch (coordinate of patch: x = 2, y = 25), run the model for 500 years and calculate the mean rate of range expansion. Previous work on the evolution of dispersal during invasions has indicated that Allee effects can play an important role [20], so we repeated the above simulations but with a mild Allee effect operating, whereby individuals that are on their own in a cell do not reproduce. We then run a set of simulations to examine how dormancy evolves during range expansion. Here, we run the simulation for 1000 years in 50 by 50 lattice to obtain evolutionary pseudo-equilibrium in a stationary environment; after this burn-in period we open-up the landscape and populations are able to expand for further 1000 years. All initial individuals are seeded with the same genotype of *γ* = 0.5. We track the eco-evolutionary dynamics of range expansion, monitoring the distribution of genotypes across space and through time, the rate of range expansion, and the abundance of adults and seeds across the landscape. We repeat the simulations for temporally stable environment and white noise, with and without a mild Allee effect operating, and for different values of *K* (5, 25) and *σ* (0.5, 1.0, 1.5, 2.0).

## Results

The rate of seed dormancy plays a major role in determining the rate of range expansion (Fig. 1). In a stable environment where seed production is constant through time, we find that higher rates of seed dormancy always result in considerably reduced rates of range expansion (Fig. 1A). If an Allee effect is operating the rate of population expansion is further reduced (Fig. 1A, blue dots). However, when seed production varies through time, intermediate rates of dormancy maximise the rate of spatial spread (Fig. 1B). Results from simulations with an Allee effect exhibit the same pattern although the overall rate of range expansion is reduced (Fig. 1B, blue dots). In a temporally variable environment, the population is prone to severe crashes (and even extinction) if dormancy is too low, especially if an Allee effect is operating (Fig. 1B).

**Figure 1.**
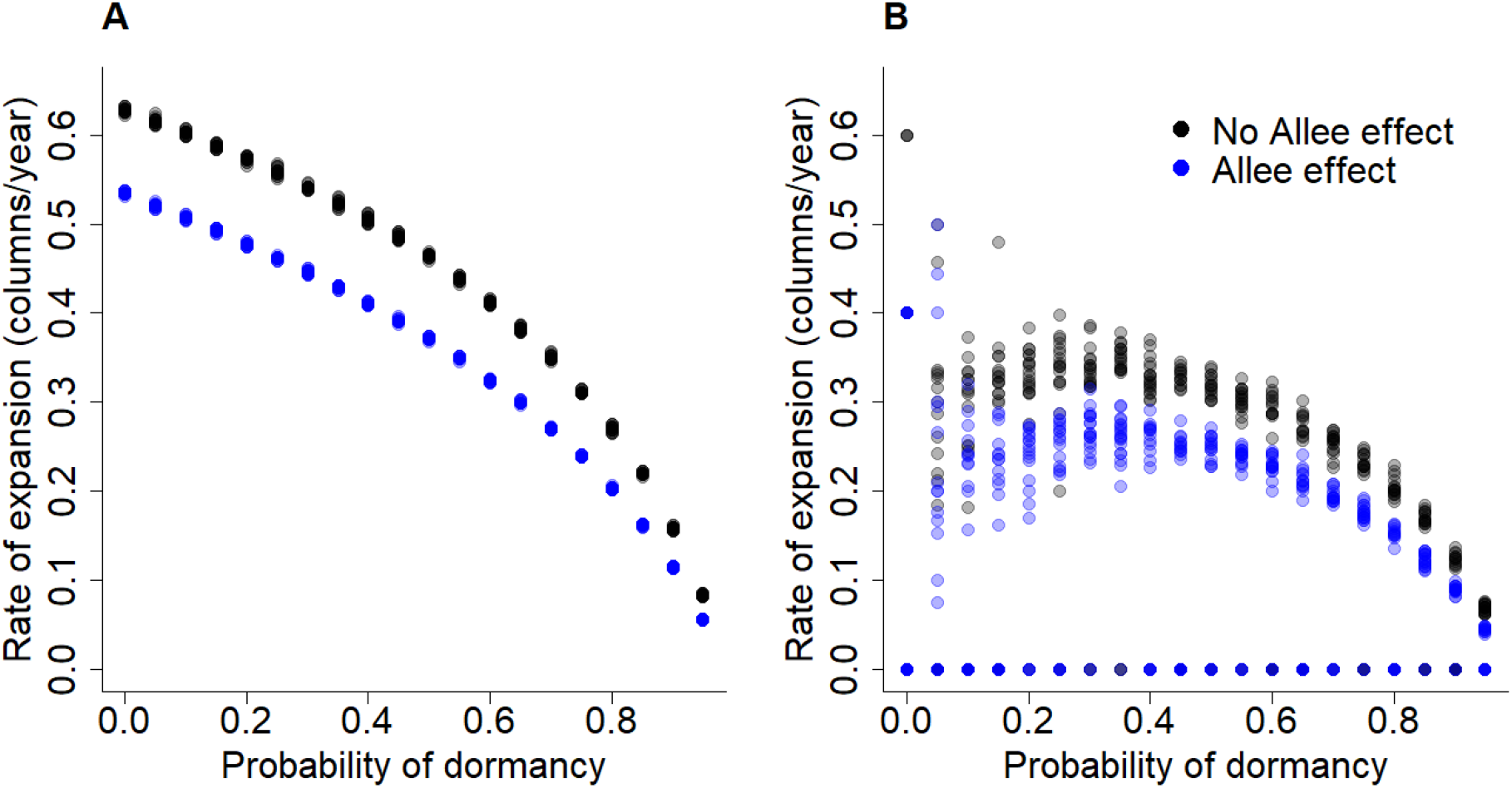
Seed dormancy controls the rate of range expansion. In a temporally stable environment **(A)** range expansion is always less rapid when the probability of seed dormancy is higher and is considerably slower when an Allee effect operates. However, in a temporally variable environment **(B)**, intermediate levels of dormancy generally result in the most rapid range expansion. Black points show the outcomes of 20 replicate simulations for each probability of dormancy without an Allee effect. Blue points show the same with an Allee effect operating. Other parameters: *d* = 0.1, *m* = 0.05, *K* = 25. In (A) *s* = 5 while in (B) *s* is subject to white noise (*κ* = 0.0, *σ* = 1.0).

Next, we consider how seed dormancy evolves during the course of range expansion. Snapshots of populations expanding their ranges into a region previously unoccupied by the species are shown in Figure 2. As the expansion proceeds, the mean rate of dormancy at the expansion front decreases, indicating that individuals with reduced dormancy are selected for. These individuals are more likely to be the first to colonise a new patch, where they are able to exploit the low intraspecific competition and thus realise higher lifetime reproductive success than conspecifics that remain dormant in the soil for some years before germinating. Behind the invasion front there is an increase in the mean rate of dormancy as the populations gradually evolve back towards the dormancy strategy that is selected in a spatially saturated environment (Fig. 2–3). Much higher dormancy evolves under white noise compared to what evolves in temporally stable environments, both before and during expansion (Fig. 3), while Allee effects do not substantially alter the evolution of dormancy. Lower local carrying capacity generally leads to evolution of higher dormancy (Fig. 3B) as kin competition is stronger in smaller populations. This effect is particularly evident in temporally stable environments, while with white noise, high environmental variability has a much bigger effect.

**Figure 2:**
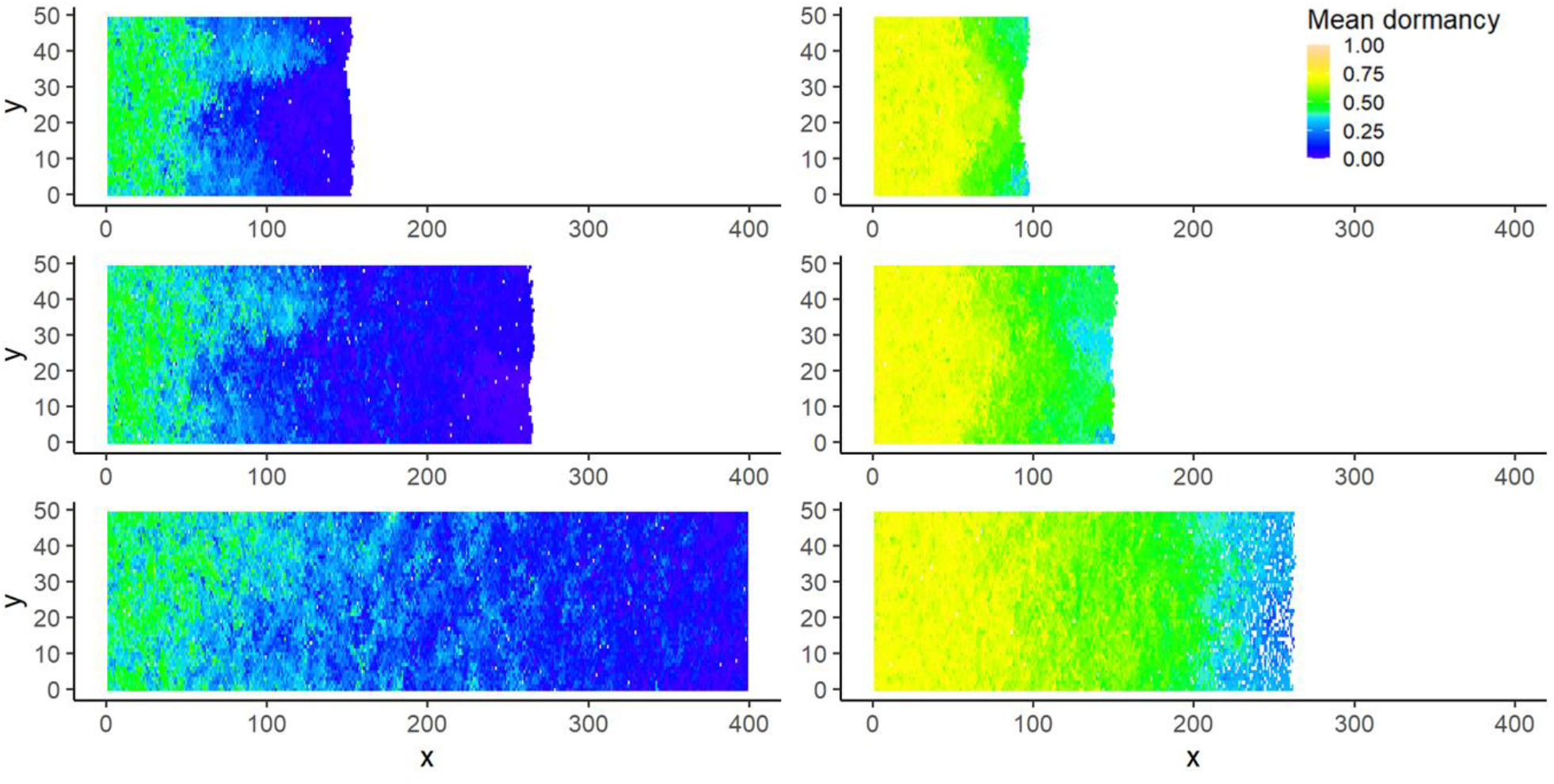
Snapshots of the model illustrating the mean probability of dormancy evolving during range expansion. The left column represents one replicate simulation under temporally stable environment, and the right column one replicate simulation under white noise, at time = 1200 (top panels), 1400 (middle) and 1775 (bottom). Both simulations are without Allee effect. Other parameters: *d* = 0.1, *m* = 0.05, *K* = 5 and *β* = 0.01.

**Figure 3:**
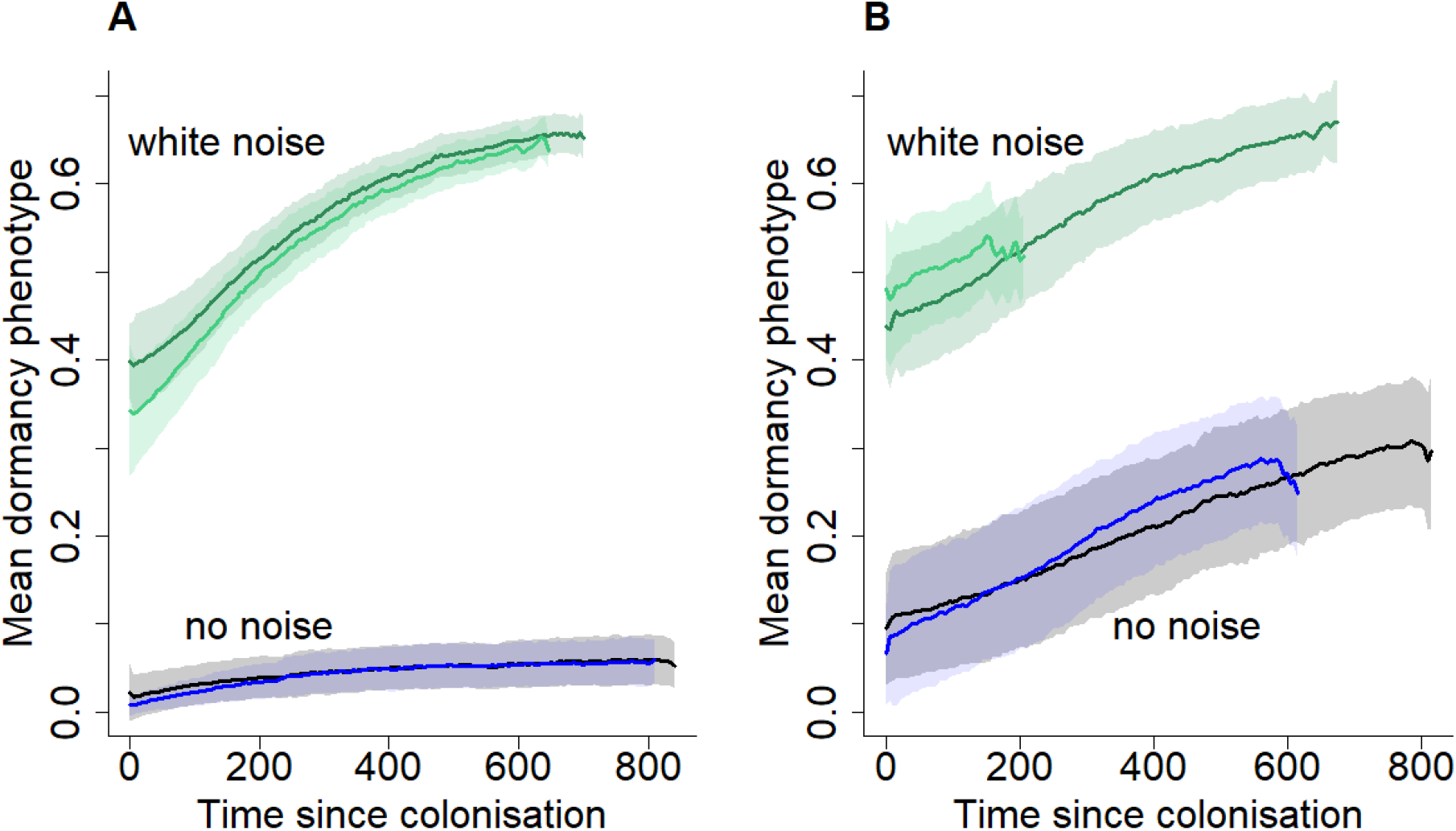
Changes in dormancy through time within a specific region, shown either under temporally stable environment or white noise (*κ* = 0.0, *σ* = 1.0), at two different carrying capacities: **(A)** *K* = 25; **(B)** *K* = 5. Here, we plot the mean dormancy of adults located in the landscape column x = 150. When the expansion front first reaches the region (time = 0) dormancy is low, but as the local population becomes established selection favours higher rates of dormancy. Dark colours (black and dark green) show simulations without an Allee effect, while lighter colours (blue and light green) show simulations with an Allee effect. Lines represent the mean dormancy phenotype averaged over 20 replicate simulations; shades represent ± the standard deviation. Other parameters: *d* = 0.1, *m* = 0.05, *β* = 0.01.

The amplitude of environmental noise (*σ*) affects the rate of dormancy that is selected both at the core and front of the range (Fig. 4). At the range core, lower amplitude fluctuations in seed production (with infrequent years of zero production) results in relatively lower dormancy evolving than for higher amplitude fluctuations which lead to higher frequencies of years with zero production. At the range front, dormancy consistently evolves to lower rates that in the core and in general there is lower dormancy at the front with lower amplitude fluctuations. However, the lowest dormancy evolves for *σ* = 1, rather *σ* = 0.5. This is likely due to the simulations with *σ* = 0.5 having reached the end of the landscape before the end of the simulation such that selection was already favouring a return towards the stationary strategy.

**Figure 4:**
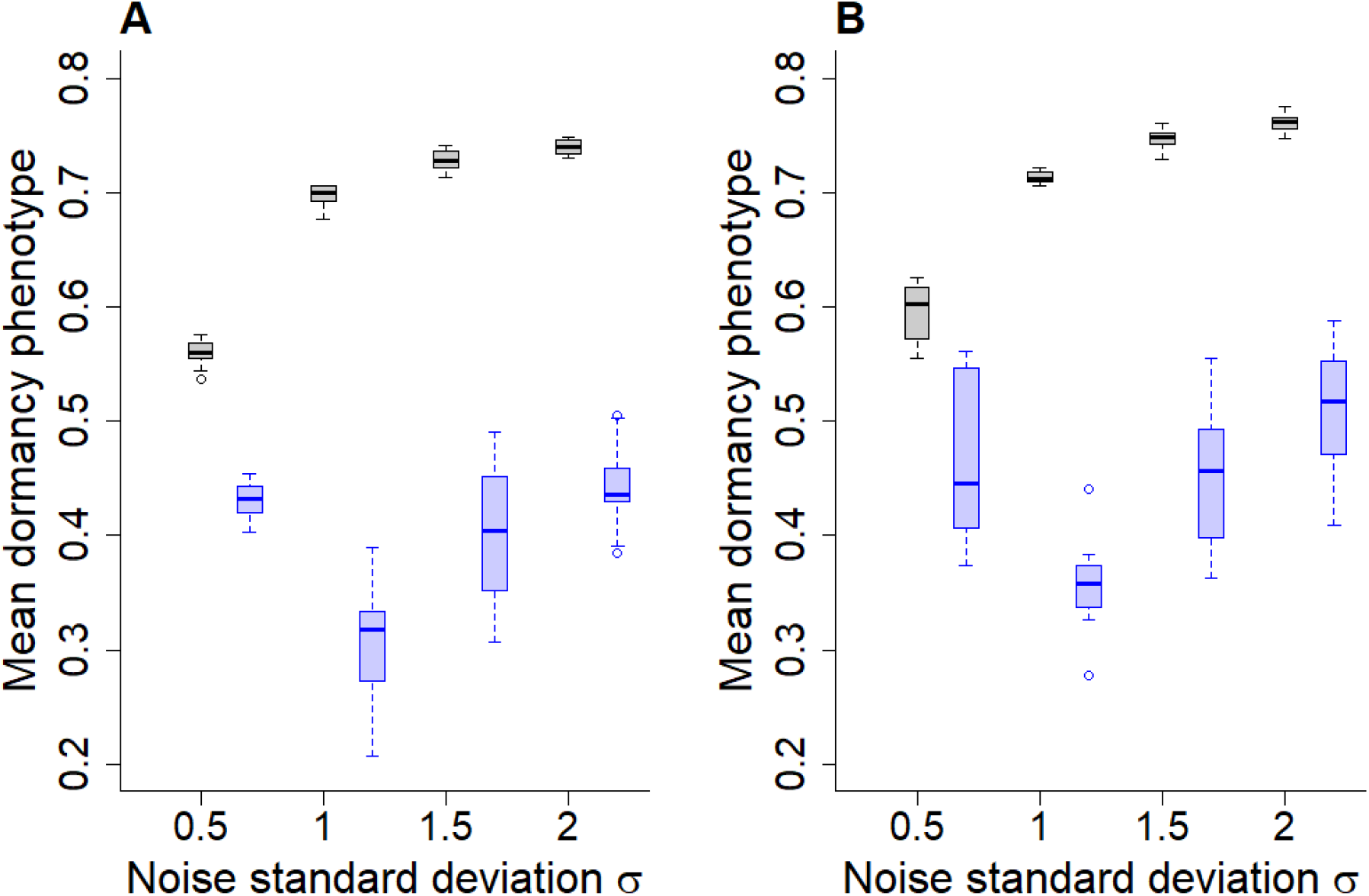
Evolution of dormancy at the core and front of the range under different amplitudes of temporal stochasticity in seed production (*σ*). **(A)** *K* = 25; **(B)** *K* = 5. Black boxes represent the mean dormancy phenotype evolved at the range core (x = 26 to 30), while blue boxes represent the mean strategy evolved in the 5 frontmost rows by year 1995. Simulations are run with no Allee effects. Other parameters: *d* = 0.1, *m* = 0.05, *β* = 0.01.

The evolution of decreased dormancy during range expansion can reduce the resilience of regional populations to adverse environmental conditions. This is neatly illustrated in Figure 5. Here, we show the abundance of adults in two equally sized regions, one that is close to the origin of the invasion and one much further away. Population abundance in the region close to the origin remains relatively stable throughout time, with just an occasional decline following a succession of poor years. The adult abundance of the more distant region reaches that of the first region roughly 80 years after the front first reaches the region. However, when the populations experience a series of poor years there is a major difference in the response of the two populations, with the local population further from the origin decreasing to much lower numbers (Fig. 5), and taking far longer to recover. Interestingly, while this difference in population resilience declines with time, there is still a detectable signal in the trajectories even after hundreds of years after the initial colonisation of the second region.

**Figure 5:**
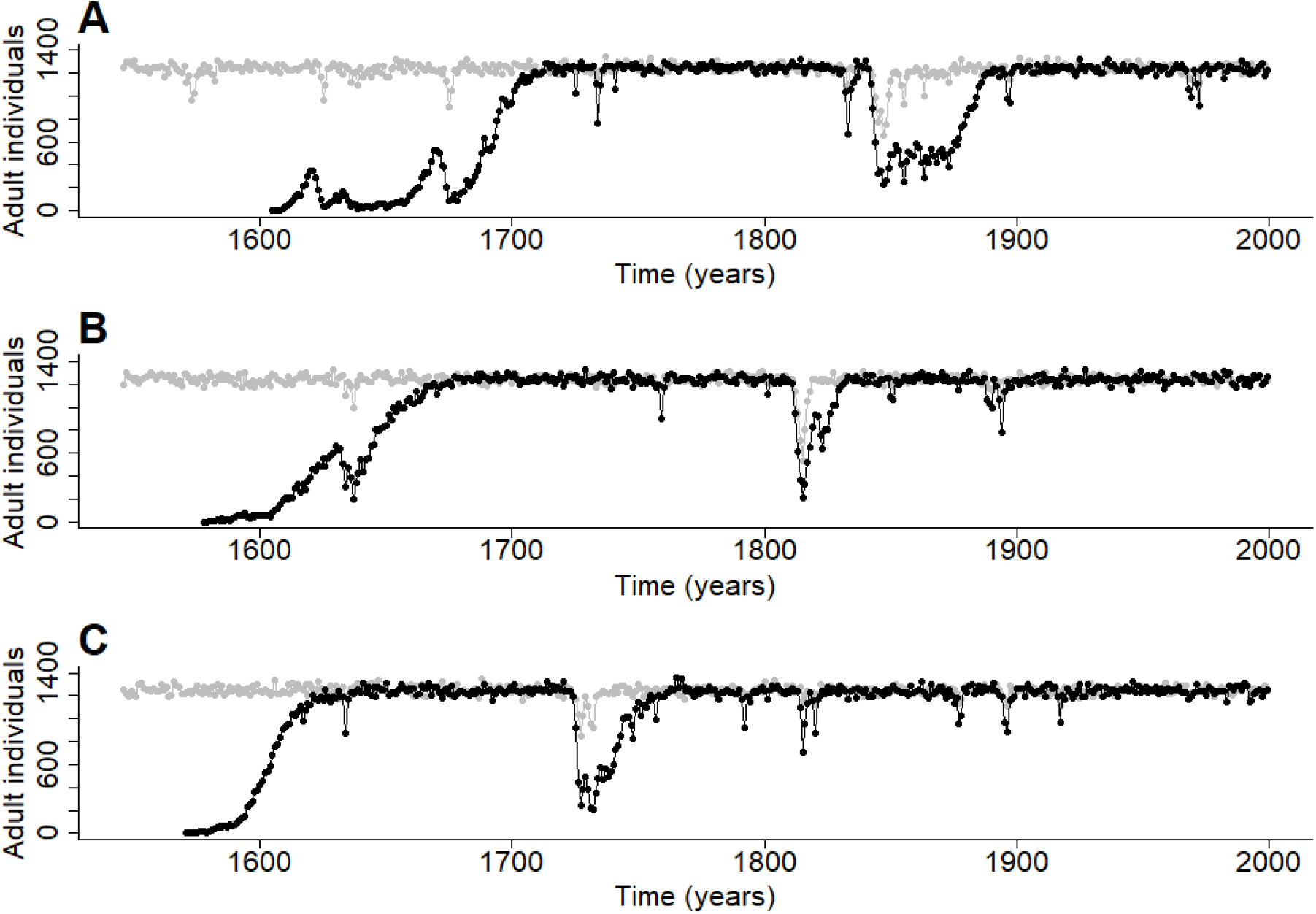
Resilience to adverse environmental conditions is decreased by the evolution of reduced dormancy during expansion. These traces (from three illustrative replicate simulations) compare the abundance of adult plants in two regions, one close to the point of introduction (x = 26 to 30; grey), the other much further away (x = 121 to 125; black), which is reached well through the expansion. These simulations are all run under white noise (*κ* = 0.0, *σ* = 1.0) and with an Allee effect. Other parameters: *d* = 0.1, *m* = 0.05, *K* = 5 and *β* = 0.01

## Discussion

Considerable evidence for the evolutionary influence on the dynamics of range expanding populations has accumulated over the last couple of decades (e.g., [2–4,6,7]). While there has been substantial work on how life history traits including dispersal and mating systems evolve at expanding fronts, dormancy has been largely neglected. Here, we have demonstrated the potential role that dormancy, and its evolution, can play in determining the dynamics of range expansions. First, we have shown that the rate of propagule dormancy controls the rate of range expansion in both temporally stable and unstable environments: in stable environments the most rapid range expansion is obtained in the absence of dormancy while in a temporally variable environment, intermediate rates of dormancy achieve the fastest rate of spread. Second, we have shown that during range expansion, selection will favour reduced rates of dormancy at the front. Third, we demonstrate that the gradient in dormancy that develops from the expanding front back towards to the core results in strong differences in the degree to which regional populations are resilient to a series of poor years.

Reduced dormancy evolves at an expanding front for similar reasons to those that lead to increased dispersal at an expanding front. Increased dispersal is selected for as it is those individuals with greater dispersal propensity or dispersal ability phenotypes that are most likely to colonise new patches at the expanding front first and, as long as there is a heritable component to the dispersal traits, their offspring will also be strong dispersers and have a high likelihood of themselves being colonists of new territory [20]. There is effectively a spatial sorting [22] of higher dispersal phenotypes at the front and this process results in rapid acceleration of dispersal during population spread. In our model dispersal is fixed at a constant emigration probability but dormancy is labile and heritable. Now, at the expanding front there is sorting of propagules produced from low dormancy individuals. If, for example, five individual propagules simultaneously arrive at an empty patch first and one breaks dormancy immediately while the others lay dormant, it will have a much higher chance of producing propagules that themselves reach another empty patch ahead of the existing front than those which exhibited dormancy. Effectively selection is acting to promote a life-history strategy that results not in a higher rate of dispersal, or in a greater dispersal distance of individuals but, by acting to reduce dormancy, in an earlier dispersal of individuals.

Previous work has indicated that Allee effects can play a major role in determining the probability that a species becomes invasive, setting the speed of an invasion [56,57], and determining the final spatial distribution of an invasive (reviewed in [58]). Allee effects have also been shown to reduce the evolution of increased dispersal during range expansion [20]. This reduced selection for increased dispersal rates arises because individuals towards the front with a higher rate of dispersal are more likely to move into a patch where they are on their own and unable to reproduce. The results described in this paper are consistent with the findings of this previous work in demonstrating that Allee effects impact on expansion speed and, additionally, indicate that they may not have the same impact on the evolution of dormancy as they do on dispersal.

A particularly interesting result, and one with potentially substantial implications, is that the evolution of a trait in a direction that confers a population with a greater rate of range expansion can at the same time reduce the population’s resilience in the face of adverse conditions. Here, reduced dormancy is selected at the expanding front, as individuals with reduced dormancy are most likely to be the first colonists of a new area and be able to achieve high fitness there, free of intraspecific competition. However, this selection for reduced dormancy can lead to populations that are ill-equipped to survive poor years, especially when a few poor years arrive in succession. This has potential consequences in the context of conservation efforts aimed at mitigating the impacts of climate change. For example, if the aim is to reintroduce populations into marginal conditions or even to assist colonisation into newly suitable climate-space, consideration should be given to dormancy characteristics of the introduced populations. For local establishment, introductions with higher dormancy stock would be beneficial but for future spread from the point of introduction, it is likely to be better to have stock with lower dormancy. Thus, an ideal strategy may be introducing a mixture of genotypes conferring differing levels of dormancy. Consequences of the eco-evolutionary dynamics of dormancy for restoration, reintroduction and assisted colonisation are an area deserving of attention and both modelling and field experiments would be useful.

The evolution of other life-history characteristics at expanding range margins might also result in reduced population resilience. As one example, at expanding margins self-incompatibility might be selected against [59–61] reducing the Allee effect and increasing the rate of range expansion. However, loss of self-incompatibility would inevitably reduce inter-individual genetic variability and potentially make the population less resilient to either a succession of poor years or to the challenge of a disease. Either through direct ecological mechanisms or through eco-evolutionary consequences, such as a reduction in adaptive potential, life history evolution at expanding fronts may result in the populations of recently colonised regions having reduced resilience against abiotic or biotic challenges. There remains much work to be done before we have a good understanding of the interplay between evolutionary and ecological dynamics at expanding range margins, and an important element of this will be to ask how evolutionary processes will alter the susceptibility of recently expanded regional populations to crashes. In tackling this question, it will be important to recognise that changes in life history traits during a range expansion may have considerable consequences on our ability to predict population dynamics. It highlights that any forecasting of the dynamics of populations in recently colonised regions that is based on life-history parameter estimates obtained from within a long-term stable range are likely to be problematic.

There are many avenues for future work investigating the evolution of dormancy during range expansions within ecological systems and beyond. We conclude by identifying five key areas that we believe merit attention.

1. Theory examining how dormancy evolves in conjunction with other life history traits during range expansions. There has been work on how evolution should be expected to shape covariation between dormancy, dispersal and mating systems within stationary ranges (e.g., [37,38,40,42,62,63]). It would be interesting to determine how predictions are likely to change in populations that are undergoing, or have recently undergone, a range expansion. Furthermore, there is the possibility that evolution of dormancy alongside other life-history traits might result in much greater acceleration of range expansions that occurs when only one trait evolves. Additionally, it would be valuable to assess how resilient recently range-expanded populations are when multiple traits have jointly evolved.
2. Empirical work examining how dormancy traits vary from an expanding front towards an introduction point (for an invasive) or a range core (for a range-shifting native). Ideally common-garden experiments are required to remove maternal (and potential grand-maternal) effects in dormancy rates. Determining the heritability of dormancy for a broad set of species would also be valuable. Related to the first area suggested above, an obvious extension would be to describe patterns of variation in both dormancy and dispersal traits from front to core.
3. Microcosm experiments have proved extremely useful in gaining improved understanding of how dispersal evolves during range expansion (e.g., [25,26,28]). Similar experiments could readily be designed to examine dormancy evolution (and joint dormancy-dispersal evolution). *A. thaliana* would provide one ideal system for such an experiment given that it has already been effectively used to explore dispersal evolution [25] and there is excellent understanding of dormancy in the species (e.g. [64–66]).
4. Theoretical and empirical work to determine how dormancy evolution during range expansion is likely to impact other adaptive responses to environmental changes. The potential for dormancy to influence the evolution of herbicide resistance [67] and drug resistance / tolerance [68] has been recognised and recent theory demonstrates that dormancy can have major impacts on adaptive processes [47]. Thus, changes in dormancy during a range expansion are likely to result in recently colonised populations having altered abilities to adapt in response to environmental change. Studies investigating this would be worthwhile and microbial microcosms would potentially prove a fruitful model system.
5. Incorporating dormancy and its evolution in tools for forecasting species’ responses to environmental change. Process-based modelling platforms are being developed that can predict ecological and population genetic responses of how species will respond to environmental change as well as being used to inform management [69,70]. These tools incorporate dispersal to differing levels of complexity, and in some cases allow for dispersal evolution (e.g., [71]). However, they do not yet incorporate dormancy, nor the potential for it to evolve. Incorporating the eco-evolutionary dynamics of dormancy would be a vital addition for effectively forecasting the future dynamics of many species.

